# On co-activation pattern analysis and non-stationarity of resting brain activity

**DOI:** 10.1101/2021.08.30.458155

**Authors:** Teppei Matsui, Trung Quang Pham, Koji Jimura, Junichi Chikazoe

**Affiliations:** Department of Biology, Okayama University, Okayama, Japan; JST-PRESTO, Japan Science and Technology Agency, Tokyo, Japan; Section of Brain Function Information, Supportive Center for Brain Research, National Institute for Physiological Sciences, Okazaki, Japan; Keio University, Yokohama, Japan; Araya Inc., Tokyo, Japan

**Keywords:** Resting-state fMRI, Co-activation pattern analysis, Non-stationarity

## Abstract

The non-stationarity of resting-state brain activity has received increasing attention in recent years. Functional connectivity (FC) analysis with short sliding windows and coactivation pattern (CAP) analysis are two widely used methods for assessing the dynamic characteristics of brain activity observed with functional magnetic resonance imaging (fMRI). However, the statistical nature of the dynamics captured by these techniques needs to be verified. In this study, we found that the results of CAP analysis were similar for real fMRI data and simulated stationary data with matching covariance structures and spectral contents. We also found that, for both the real and simulated data, CAPs were clustered into spatially heterogeneous modules. Moreover, for each of the modules in the real data, a spatially similar module was found in the simulated data. The present results suggest that care needs to be taken when interpreting observations drawn from CAP analysis as it does not necessarily reflect non-stationarity or a mixture of states in resting brain activity.

**Highlights:**

- CAP was conducted for real fMRI data and a stationary null model.
- The results of CAP analysis were similar for the real and simulated data.
- Similar results for real and simulated data in both ROI- and voxel-based analyses.
- Results of CAP analysis may not reflect non-stationarity of resting brain activity.

## Introduction

In contrast to initial studies of resting-state functional connectivity (FC), which assumed FC was stable throughout a relatively long scan-duration, recent studies are increasingly focusing on non-stationary aspects of resting brain activity (Calhoun et al., 2014; Hutchison et al., 2013; Mitra et al., 2018; Preti et al., 2016; Shine et al., 2016). This dynamic view of resting brain activity hypothesizes that the brain switches between various states within which the activity of different sets of brain regions are coordinated. Sliding window correlation analysis and co-activation pattern (CAP) analysis have been developed to assess such non-stationary characteristics of resting brain activity (Allen et al., 2014; Karahanoğlu and Van De Ville, 2015; Liu et al., 2013; Liu and Duyn, 2013).

Sliding-window correlation analysis and CAP analysis detect momentary brain activity patterns by using FC within a short sliding-window (∼40 sec) (Allen *et al*., 2014) and brain activations in selected single volumes (Liu *et al*., 2013; Liu and Duyn, 2013), respectively. The detected momentary brain activity patterns are often heterogeneous, even within the same run, and grouped into distinct clusters of modules. In both sliding-window correlation analysis and CAP analysis, these modules exhibit specific spatial patterns which are therefore interpreted as distinct states of the resting brain (Liu and Duyn, 2013; Liu et al., 2018). Numerous studies have applied these methods to characterize the non-stationary aspects of resting brain activity in healthy individuals, patients with neurological disorders and animals [see (Liu *et al*., 2018; Preti *et al*., 2016) for review].

Importantly, whether the results obtained from sliding-window correlation analysis or CAP analysis truly reflect the non-stationarity of resting brain activity is a crucial issue that needs careful study (Handwerker et al., 2012; Hindriks et al., 2016; Liégeois et al., 2017; Lurie et al., 2020; Zalesky and Breakspear, 2015). In fact, several studies have shown that some results claimed to reflect the non-stationarity of resting brain activity can be replicated from simulated data with stationary null models (Cifer et al., 2017; Laumann et al., 2016; Novelli and Razi, 2021). The stationarity of resting brain activity implies, in contrast to the commonly held assumption, that the resting brain does not alternate between meta-stable states (Hutchison *et al*., 2013) but occupies a single resting-state. On the other hand, the non-stationarity of resting brain activity implies an alternation between meta-stable states. Each meta-stable state exhibits a functional connectivity that is distinct from other meta-stable states. [See (Liégeois *et al*., 2017) for further details of the stationarity and non-stationarity of resting-brain activity]. Although these studies do not prove or claim that resting brain activity is stationary, they provide the necessary and important background required for appropriately interpreting the results obtained with these analyses designed to capture the non-stationarity of resting brain activity. In this report, we add to this discussion by examining whether simulated stationary data can reproduce the spatially heterogeneous modules obtained with CAP analysis (Liu *et al*., 2018). To our surprise, we found that CAP analysis applied to a stationary null model yielded spatially heterogeneous modules, each of which closely approximated the modules found in the real data.

## Materials & Methods

### Dataset

We used S1200 release of resting-state fMRI distributed by the Human Connectome Project (HCP; http://humanconnectomeproject.org/)(Van Essen et al., 2013). The data was a collection of region-of-interest (ROI) time series (1200 volumes × 264 ROIs × 64 individuals; Repetition time (TR), 0.72 sec) (Power et al., 2011). Note that the data went through several preprocessing including ICA-based denoising (ICA-FIX) (Glasser et al., 2013). Each time course was normalized to have zero mean and unit variance. Global signal regression was conducted by following a standard procedure. For each scan, a global signal time course was obtained by averaging time courses across all ROIs. Then for each ROI, the global signal time course was regressed out from the ROI’s time course. The data for group analysis was made by concatenating data of all the individuals in the volume dimension (Liu and Duyn, 2013).

For voxel-based analyses, we selected a single axial slice (z = 27 in the MNI coordinate) containing the posterior parietal cortex (PPC) in the HCP resting-state data to mitigate computational demand. The data of 60 individuals were used in the analysis. All data were preprocessed according to the HCP’s standard pipeline. To select the voxels corresponding gray matter, we made a gray matter mask for each individual using the segmentation program implemented in SPM12 (https://www.fil.ion.ucl.ac.uk/spm/software/spm12/; threshold set at 0.7) and then took a union of the masks. The obtained mask was applied to each axial slice to extract 2320 voxels corresponding to the gray matter.

### Static FC

Static FC was calculated by taking the correlation coefficient between the time courses of two ROIs using all volumes in the concatenated runs (Fox et al., 2005; Matsui et al., 2011). For the voxel-based analyses, an ROI centered at the PPC (6 mm by 6 mm centered at MNI coordinates [0, -53, 27]) was used, and the temporal correlation was calculated between the ROI and each voxel.

### Generation of simulated data

The simulated data were generated by adapting previously described methods and codes (Laumann *et al*., 2016; Matsui et al., 2018). Briefly, random samples were drawn from a Gaussian distribution with dimensions matching the real data. These time courses were multiplied in the spectral domain by the spectrum derived from the real data, ensuring that the power spectra of real data were preserved. The time courses were then projected onto eigenvectors of the covariance matrix derived from the real data, ensuring that the covariance structure of the real data was preserved. This procedure enabled the construction of simulated time courses, stationary by construction, with covariance structures and spectral contents equivalent to the real data. Here after, we call this null model the Laumann null. Unless otherwise noted, the Laumann null was used to produce simulated data. Simulated time courses were generated for each run and each individual. The results were similar when the data were generated for the real data concatenated across individuals (data not shown). Global signal regression was similarly performed for the real data. Group data were then obtained by concatenating all the simulated data across individuals.

In addition to the Laumann null, three null models were additionally tested. The first was a static null, which retained only the covariance structure of the real data (static null). Simulated data for static null were generated using a multivariate Gaussian whose covariance matrix was set to the covariance matrix of the real data. The second was an autoregressive randomization null model (ARR null). The lag of ARR null was set to 1. Thus, ARR null assumed that the fMRI data at time *t* is a sum of the linear transformation (*A*_*1*_) of the fMRI data at time *t-1* and a zero-mean multivariate Gaussian noise with covariance matrix (*Σ*). The parameters for the autoregressive equation (*Σ,A*_*1*_) are fitted as described previously (Liégeois *et al*., 2017). Simulated data for ARR null were generated using a randomly selected time point from the real fMRI data as the seed and by iteratively applying the autoregressive equation. The third model was a phase randomization null model (PR null). PR null retained the complete autoregressive structure of the real data as well as the covariance structure (Liégeois *et al*., 2017). Simulated data for PR null were generated by first applying discrete Fourier transform (DFT) to the real fMRI data. Random phases were added to the Fourier transformed data, and then inverse DFT was applied. Added phases were independently generated for each frequency, but were the same across brain regions (Liégeois *et al*., 2017). Note that static null and Laumann null did not retain cross-spectral properties (unlike ARR null and PR null).

### CAP analysis

The CAP analysis was conducted according to standard procedures (Liu and Duyn, 2013; Liu *et al*., 2018). For each ROI time course, time points exceeding the percentile threshold (top 15%, unless otherwise noted) were collected. The set of volumes corresponding to these time-points were defined as CAPs (using the ROI as Seed). In the group analysis, for each chosen ROI, CAPs were selected using the concatenated time courses. For the voxel-based analyses, the same PPC ROI used to calculate the static FC was used. Average CAP (for a Seed ROI) was obtained by averaging across all detected CAPs. Modules were extracted by k-means clustering of the CAPs using the correlation distance. The number of clusters was set to eight as in the original study (Liu and Duyn, 2013). The similarity of the modules, both within and across data types, was measured by calculating the ROI-wise or voxel-wise correlation between two modules. The distribution of states was calculated as the fraction of each module in the total number of CAPs. The transition probability was calculated as the probability that module A was followed by module B in the consecutive CAPs. The transition probability matrix was obtained by calculating the transition probability for all combinations of modules. In the transition probability matrix, the probability was normalized for each seed module (*i*.*e*. module A). Comparisons of the real and the simulated distributions of state were done by calculating the correlation of the two after reordering the modules of the simulated data (to maximize the match with the real modules). Similarly, comparisons of the real and the simulated transition probability matrices were done by calculating the elementwise correlation of the two after reordering the modules of the simulated data.

### Statistical comparison of the real and simulated CAPs

For each pair of real and simulated CAPs obtained with a seed ROI, we tested the null hypothesis that the two sets of random multivariate variables, *i*.*e*., the real and simulated CAPs, were drawn from the same distribution using energy statistics (Szekely and Rizzo, 2013). This statistics quantifies statistical distance between the distributions of random vectors, which characterizes equality of the distributions. We used a Matlab implementation provided by Dr. Brian Lau to analyze the energy statistics. (R codes developed by the original authors are also available [https://github.com/mariarizzo/energy]). Because of a limitation in computational power, the real and simulated CAPs were each subsampled to 1000 samples before being subjected to the energy test. Covariance matrices were compared by taking the correlation coefficient of the off-diagonal elements.

### Data and code availability

All data used in the present study are from distributed by HCP. A code for reproducing essential results is available for download(https://github.com/teppei-matsui/CAP). All codes used for the analysis will be provided upon reasonable request to the corresponding author.

## Results

### A small number of volumes suffices to approximate static FC in both real and simulated data

A key observation in the CAP analysis was that static FC could be closely approximated with a small fraction of time points. Because this observation was the conceptual starting point of the CAP analysis (Liu and Duyn, 2013), we first examined if this property was genuine to the real fMRI data or reproducible with the simulated stationary data. Figure 1 shows comparisons between static FC, the average CAP of the real data, and the average CAP of the simulated data whose covariance structure and spectral contents were matched to the real data. Supplementary Figure 1 shows similar comparisons for two representative individuals. These results suggest that a small number of volumes can be used to approximate static FC for both real and simulated data. Consistent results were found for CAPs obtained using the top 5% as the threshold (Supplementary Figure 2) and for the analysis without global signal regression (Supplementary Figure 3). Thus, these results suggest that the stationary null model replicated the observation that a small fraction of time points was sufficient to approximate static FC.

**Figure 1.**
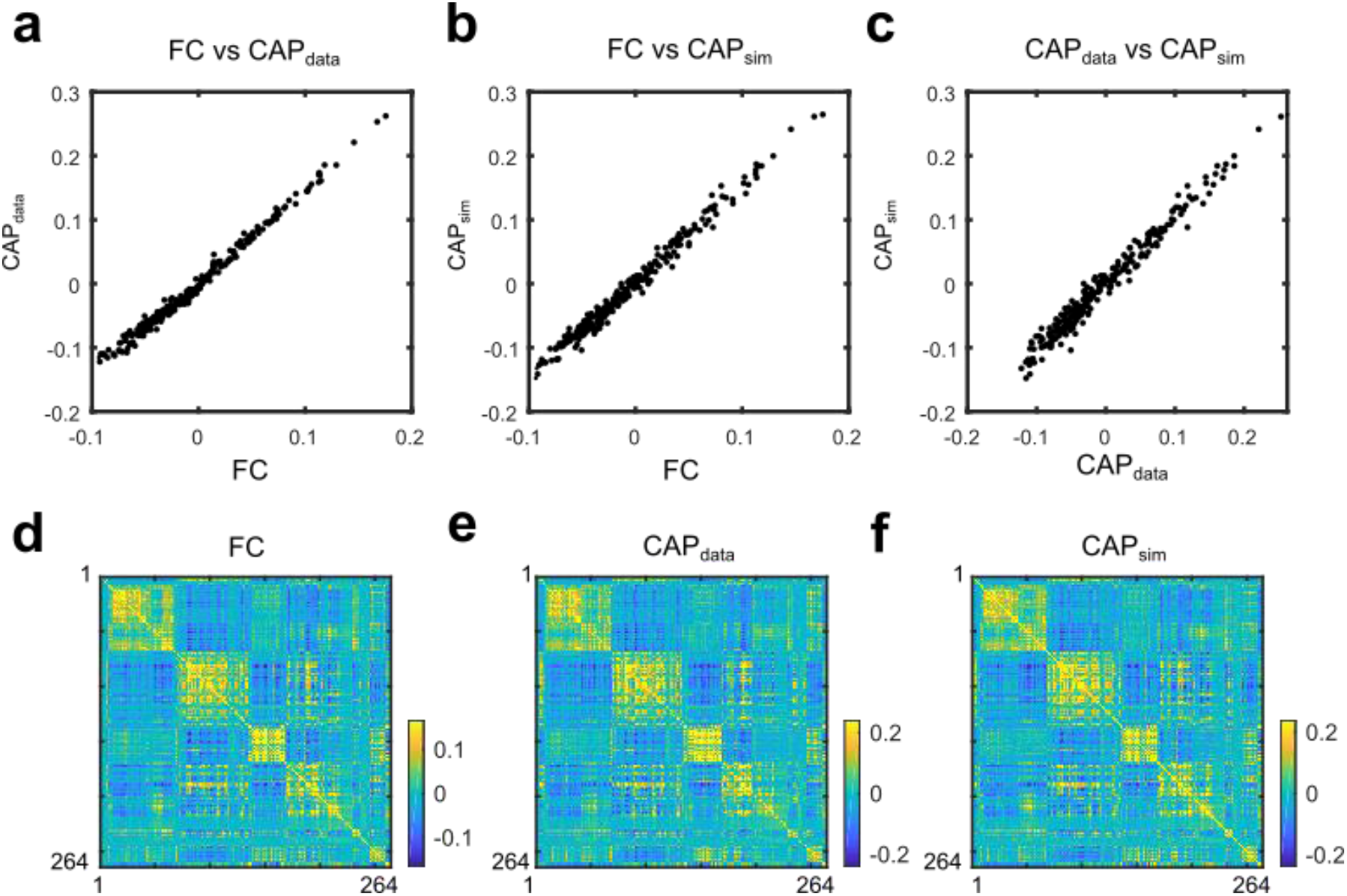
The average CAPs of the real and simulated data closely approximated static FC. **a-c**. Scatter plots of a representative ROI showing close correspondence between FC, CAP_data_, and CAP_sim_. **d-e**. ROI by ROI matrices of FC, CAP_data_, and CAP_sim_.

### Both real and simulated CAPs can be clustered into heterogeneous modules

Another key finding from the initial CAP analysis was that CAPs could be clustered into modules with distinct spatial patterns. The presence of these modules was interpreted as distinct states of resting-state brain dynamics and regarded them as an indication of non-stationarity (Liu and Duyn, 2013). If this interpretation were true, CAPs obtained in the stationary null model would not yield spatially heterogeneous modules. To test this, we clustered CAPs into modules for both the real and simulated data. Figure 2 shows the similarity between modules, within the same data type, for the real and simulated data. The clustering of CAPs resulted in spatially heterogeneous modules (*i*.*e*., low spatial correlation between modules) for both real and simulated data. Thus, the presence of spatially heterogeneous modules cannot be taken as evidence for non-stationarity in resting brain activity.

**Figure 2.**
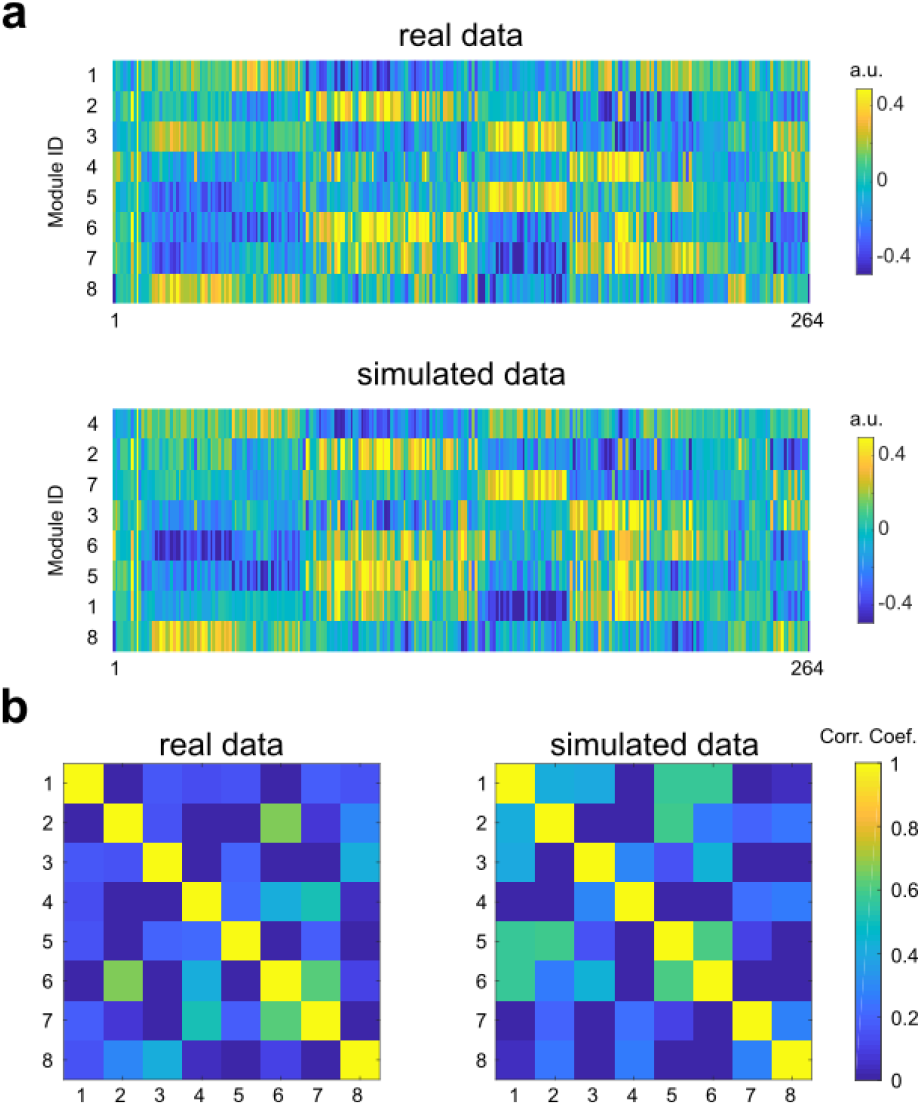
Spatially heterogeneous modules were found both for the real and simulated CAPs. **a**. Example of CAP modules for the real (top) and simulated data (bottom). Module IDs of the simulated data were reordered to maximize the match with the real CAP modules (see also Figure 3). **b**. Colored matrices show similarities between modules obtained by clustering the real or simulated CAPs. Left, real data. Right, simulated data.

This result seemingly contradicts the original report by Liu and Duyn where they found no modular structure in CAPs obtained with control (simulated) data generated using normal distributions (Liu and Duyn, 2013). Importantly, unlike the simulation devised by Laumann and colleagues (Laumann *et al*., 2016), the simulation used by Liu and Duyn did not impose spatial covariance among voxels (*i*.*e*., each voxel treated as independent). We found that simulated data with zero spatial covariance among ROIs greatly increased the similarity between CAP modules (Supplementary Figure 4), replicating the results described by the previous study. Thus, the apparent contradiction between the present simulation results (Figure 2) and Liu and Duyn’s study is due to the difference in null model construction. Because the assumption of zero spatial covariance among voxels or ROIs is unlikely to hold for real brain activity, we only considered the Laumann-type null model in the following analyses.

Having seen that the stationary null model can produce heterogeneous modules of CAPs, we next proceeded to ask whether modules of CAPs obtained from the real and simulated data were similar. If the real and simulated modules were alike, it would mean that the spatial pattern of modules is primarily determined by static properties of the real data, *i*.*e*., the covariance structure and spectral contents. Figure 3 shows (between datasets) a comparison of the modules obtained from the real and simulated data. Unexpectedly, for each module obtained from the real data, a similar module could be found from the simulated data (Figure 3a). To quantify the similarity of the two sets of modules, we matched the two sets to maximize the mean of the diagonal elements of the correlation matrix (Figure 3b). The similarity index defined by the mean of the diagonal elements was 0.83. Importantly, this value was within the range of the similarity index calculated by comparing two sets of modules derived from two independent simulations (mean ± SD, 0.81 ± 0.045, n = 100 pairs of simulations; Figure 3c), indicating that the similarity between the real and simulated modules was close to the noise ceiling set by statistical sampling error. A high degree of similarity was found between the real and simulated modules across all ROIs (mean ± SD, 0.81 ± 0.037, n = 264 ROIs; Figure 3d). Taken together, these results suggest that CAPs were clustered into spatially heterogeneous modules in both real and simulated data. Moreover, individual modules obtained from the simulated data were similar to modules obtained from the real data, suggesting that the spatial pattern of modules is largely determined by static properties of resting brain activity.

**Figure 3.**
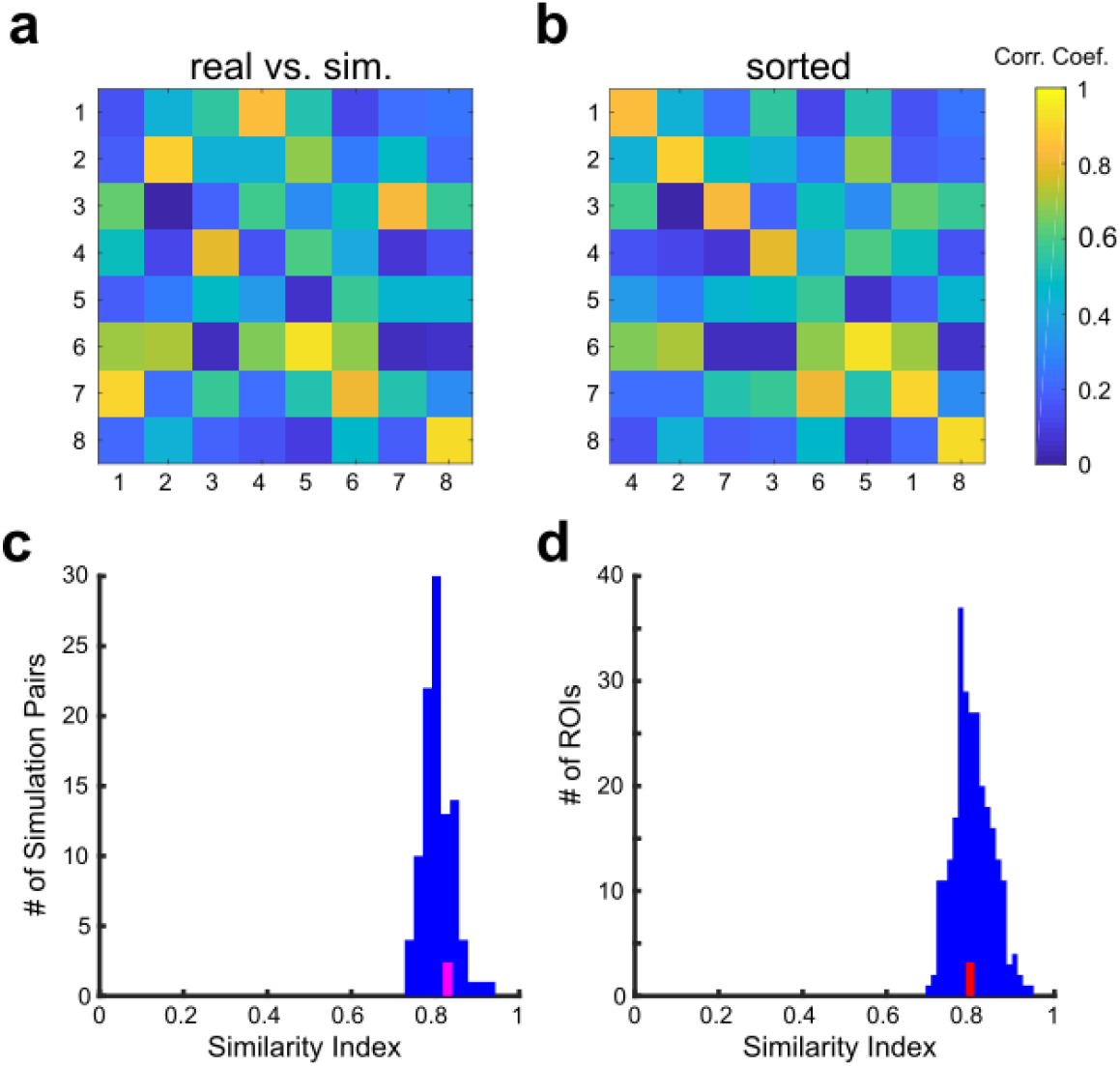
CAP modules derived from real and simulated data were very similar. The similarity between modules was measured by the correlation coefficient. **a**. original ordering as in Figure 2. **b**. same as *a* but the modules were reordered to maximize matching (mean of diagonal elements = 0.83). **c**. similarity distribution of modules between two independent simulations (same ROI as in *a,b*). For each pair of simulations, module matching was performed (as in *b*), and the mean of the diagonal elements was taken as the similarity value. The magenta line indicates the similarity value for the data sown in *a,b*. **d**. similarity distribution of the modules between the real and simulated data tested for all ROIs. The magenta line indicates the similarity value for the data sown in *a,b*.

To specify further the statistical properties of the real data that produced these results, we tested three additional null models (see Methods). The first null model retained only the covariance structure of the data (static null). The second null model was an autoregressive model with lag equal to 1 (ARR null). The last null model was a phase randomized data (PR null). The results obtained with the Laumann null were reproduced with all three null models (Supplementary Figures 5). The similarity index of CAP modules was 0.81 ± 0.043 for static null, 0.80 ± 0.040 for ARR null, and 0.81 ± 0.044 for PR null (mean ± SD, n = 264 ROIs; recall that the value for the Laumann null was 0.81 ± 0.047). The fact that the results obtained with the static null did not differ from the results obtained with the other null models suggests that the spatial structures of CAP and modules were determined by the covariance structure of the real data.

We also examined the distributions of states (i.e. CAP modules) and matrices of transition probability between states to analyze the temporal structure of CAPs. We found that the distributions of states were similar for real data and the four null models (Supplementary Figure 6a). The correlation of the distribution of the state in real and that of a null model was 0.56 ± 0.27 for the Laumann null (mean ± SD., n = 264 ROIs), 0.53 ± 0.28 for static null, 0.53 ± 0.29 for ARR null and 0.56 ± 0.26 for PR null (Supplementary Figure 6c). The transition probability matrices differed among null models (Supplementary Figure 6b): The matrix for static null appeared very different from that of the real data. The matrix for the Laumann null was more similar to that of the real data than static null but was less so than ARR null and PR null. The matrices for ARR null and PR null were similar. The element-wise correlation of the transition probability matrix in the real data and that of a null model was 0.84 ± 0.040 for the Laumann null, 0.38 ± 0.12 for static null, 0.96 ± 0.017 for ARR null, and 0.95 ± 0.018 for PR null (Supplementary Figure 6d). High values of the element-wise correlation in ARR null and PR null were consistent with a previous report that these models closely recapitulate the dynamics of resting fMRI activity (Liégeois *et al*., 2017). The fact that ARR null and PR null better reproduced the dynamics of real CPAs than the Laumann null suggests that between (auto- and cross-) covariance with lag 1 contributed significantly to the dynamics of CAPs.

We next conducted analyses with voxel-based data to visualize each module and confirm that the spatial averaging for the ROI-based analysis did not cause the similarity between real and simulated data. To mitigate computational demand, we performed this analysis on a single slice containing PPC (see Methods). Overall, the results from the voxel-based analysis were similar to those of the ROI-based analysis described above. Like the results shown in Figure 1a-c, we found the average CAP_data_ and CAP_sim_ maps were similar to the static FC map (Figure 4a). For both the real and simulated data, modules of CAPs were dissimilar to each other within the same data type (Figure 4b,c; see also Figure 2 for comparable results in the ROI-based analysis). Finally and most importantly, simulated modules with similar spatial patterns were found for most of the modules in the real data (Figure 4d,e). Figure 5 shows spatial maps of the modules shown in Figure 4d,e. These examples visually demonstrate the high degree of similarity between the real and simulated modules. Thus, the voxel-based results corroborate the findings of the ROI-based analyses and further show that the presence or absence of spatial averaging does not change the results.

**Figure 4.**
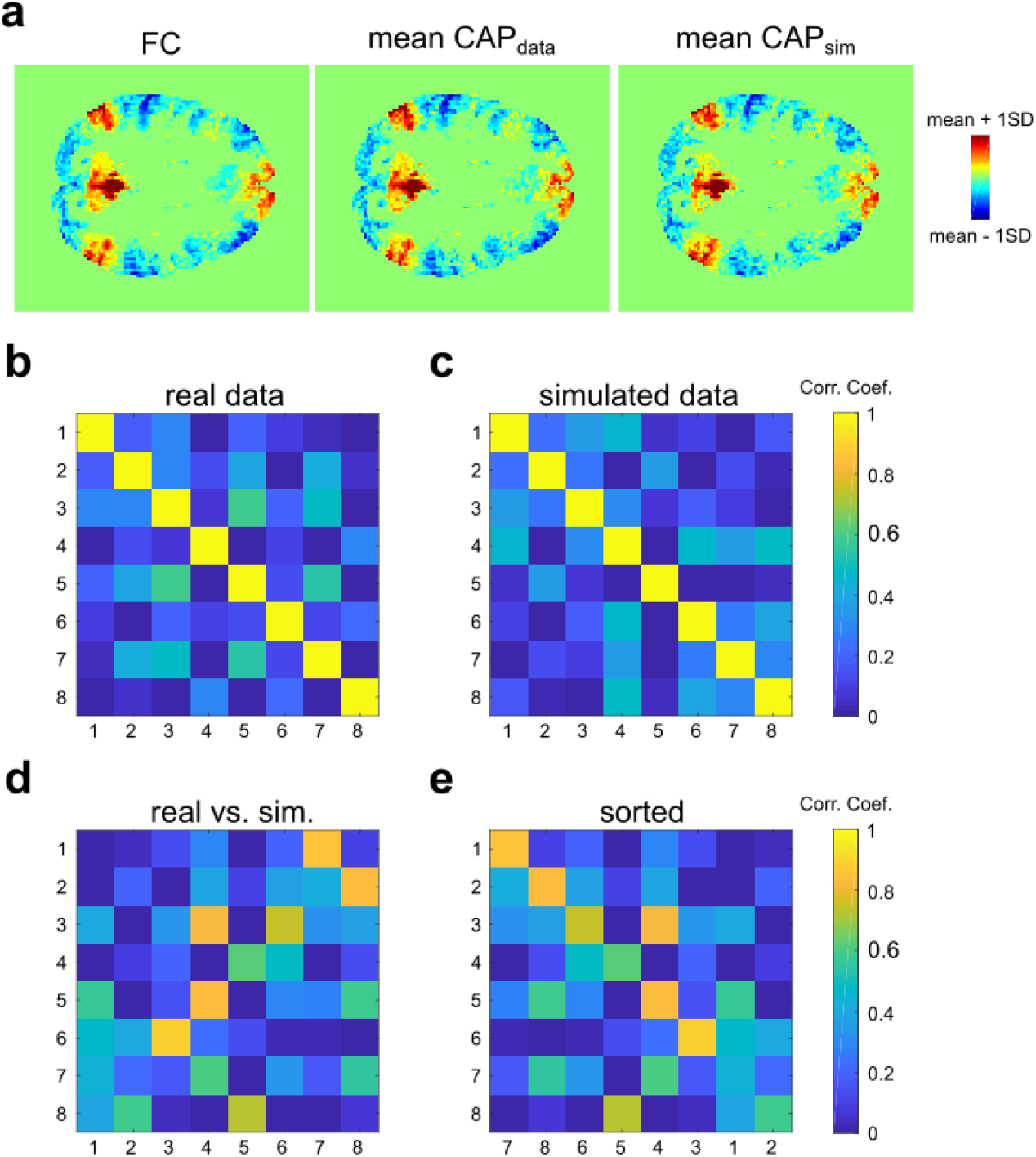
Voxel-based analysis yielded the similar results to the ROI-based analysis. **a**. Comparison of the FC map, mean CAP_data_ map, and mean CAP_sim_ map. As in Figure 1, the three maps appear very similar. **b-c**. The similarity between CAP modules within the same data type. Following the same convention as in Figure 2. **d-e**. The similarity between modules across data types. **d**. original ordering as in (b)-(c). **e**. Same as (d) but modules are reordered to maximize matching (mean of diagonal elements = 0.72). See Figure 5 for details of each module.

**Figure 5.**
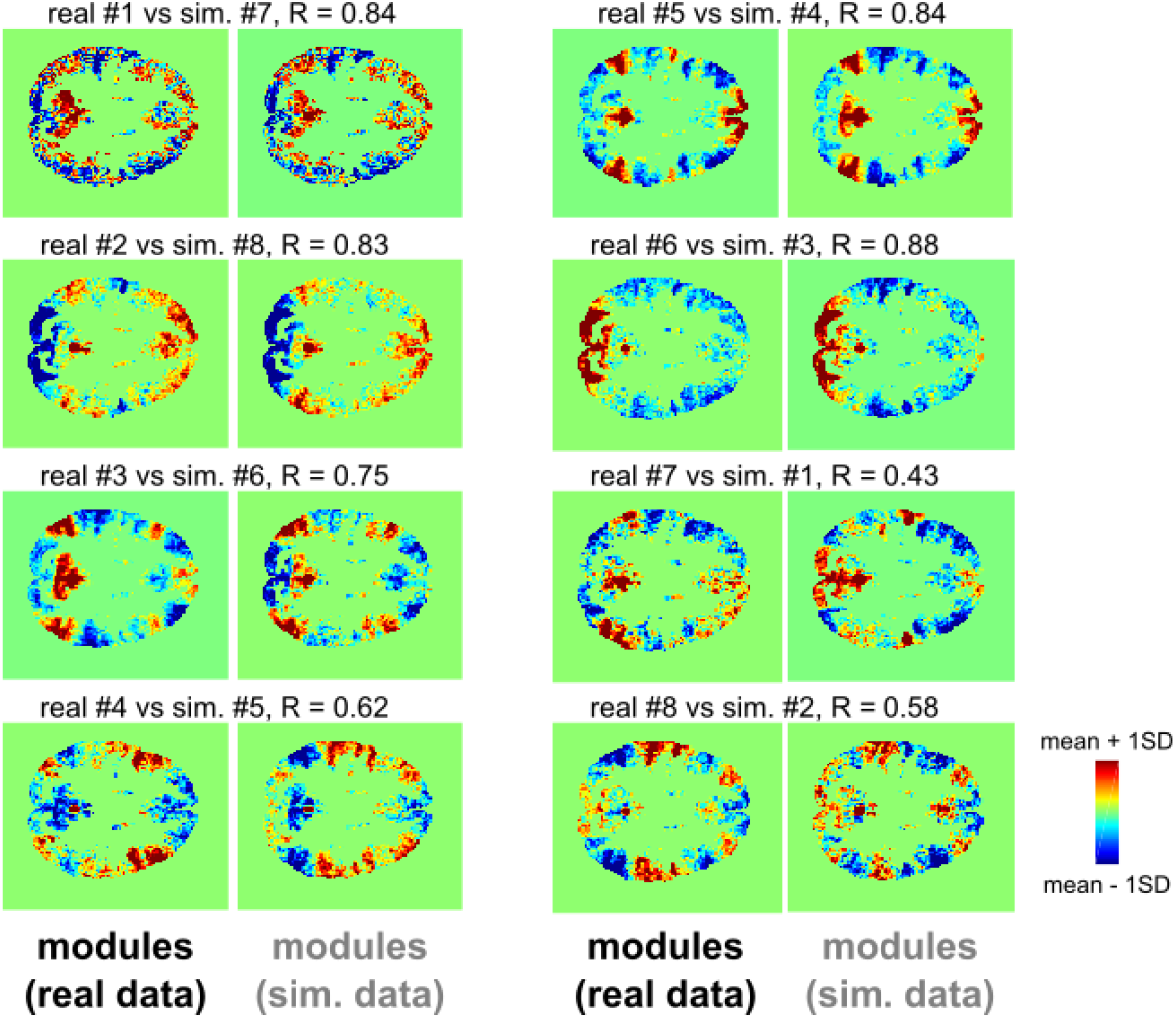
Examples modules derived from the real and simulated data. For the example shown in Figure 4, eight pairs of modules derived from the real data (left) and the simulated data (right) are shown side-by-side. The modules are paired according to Figure 4e.

Why were modules of CAPs in the real data so similar to those in the simulated data? We hypothesized that this observation was because the real and simulated CAPs shared the same underlying statistical distribution. In k-means clustering, data points were grouped into clusters according to the correlation distance between the data points. Evidently, the distribution of the correlation distance was determined by the distribution of the data points. Therefore, if the datasets were derived from the same distribution, provided enough data points were included in each set, the results of k-means clustering should have been similar.

To compare the statistical similarity of the two sets of CAPs, we first compared the covariance matrices. Note that the covariance matrices were calculated only using CAPs and were hence different from those using all volumes. For a representative ROI, we found the covariance matrices of the real and simulated CAPs were very similar (R = 0.97; Figure 6a,b). A high degree of similarity was found between covariance matrices of the real and simulated CAPs for all ROIs (mean ± SD, 0.97 ± 0.0026, n = 264 ROIs; Figure 6c), consistent with the hypothesis that they share the same multivariate distribution. Next, we conducted statistical testing to examine the null hypothesis that the two datasets were drawn from the same multivariate distribution (see Methods). If the real and simulated CAPs were derived from different distributions, the null hypothesis should be rejected. Statistical testing was not significant for the representative ROI used in Figure 6a (P > 0.87). Across all ROIs, p-values were mostly distributed above the typical significance threshold (mean ± SD, 0.49 ± 0.30; Figure 6d; Note that p-values were uncorrected). The null hypothesis was rejected in 20 out of 264 ROIs (7.6% of all ROIs) when the significance threshold was 5%. With a significance threshold of 1%, the null hypothesis was rejected in 7 ROIs (2.7% of all ROIs). Thus, for most ROIs, the statistical distributions of the real and simulated CAPs were similar and hence yielded similar clustering results.

**Figure 6.**
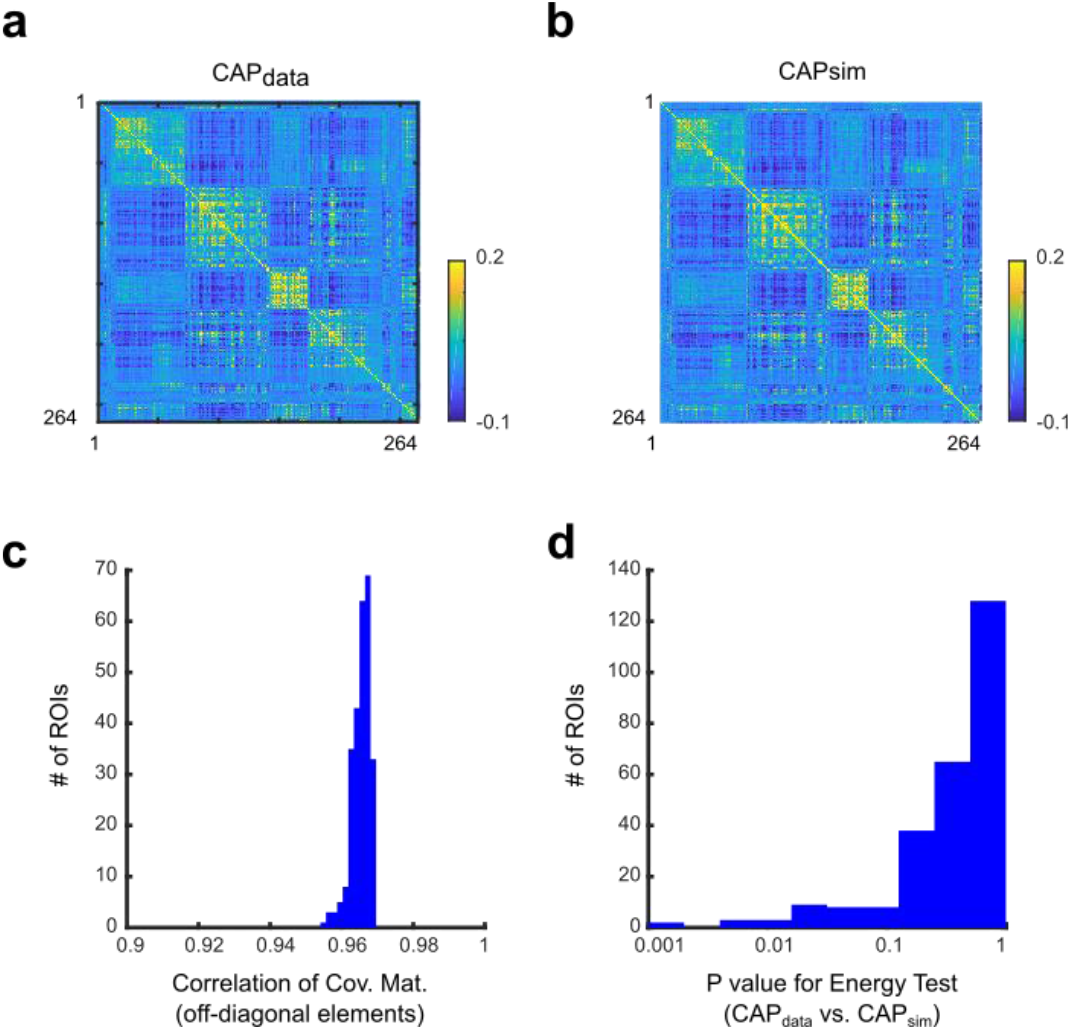
Statistical properties of the real and simulated CAPs were approximately equal. **a-b**. ROI-by-ROI covariance matrices of the CAP_data_(a) and CAP_sim_(b). **c**. The correlation distribution between off-diagonal elements of the covariance matrices of the real and simulated CAPs (n = 264 ROIs). **d**. The distribution of p-values from Szekeley & Rizzo’s Energy Test, which tests the null hypothesis that CAP_data_ and CAP_sim_ are drawn from the same multivariate distribution (One test per ROI. n = 264 ROIs).

## Discussion

In this study, we asked to what extent could the results of CAP analysis be attributed to the non-stationarity of resting brain activity. To this end, we conducted CAP analysis on real fMRI data and simulated data based on a stationary null model with matching covariance structures and spectral contents. Overall, we found that two key observations drawn from the CAP analysis were replicated in the simulated data, which has previously been interpreted as evidences for non-stationary resting brain activity. First, in both the real and simulated data, a small percentage of time points were sufficient to approximate FC calculated using all time points (Figure 1 and Figure 4a). Second, both the real and simulated CAPs were classified into spatially dissimilar modules (Figure 2 and Figure 4b,c). More interestingly, it was possible to find a simulated module closely resembling the spatial pattern of most modules obtained from the real data (Figure 3, Figure 4d,e and Figure 5). The fact that key results of CAP analysis were replicated in simulated stationary data suggests that the results need to be interpreted with care.

The present study adds to a series of previous studies reporting that it is difficult to find signatures of the non-stationarity of resting brain activity in fMRI data (Cifer *et al*., 2017; Hindriks *et al*., 2016; Laumann *et al*., 2016; Novelli and Razi, 2021). The temporal variability in FC observed with sliding-window FC analysis has been attributed to statistical sampling error (Hindriks *et al*., 2016; Laumann *et al*., 2016). Notably, Laumann and colleagues developed a stationary null model based on a multivariate Gaussian distribution and matching covariance structures and spectral contents to real data. They used this null model to show that results obtained with sliding window FC analysis are similar for real and simulated data, suggesting that the stationarity of the data cannot be distinguished based on sliding window FC analysis. In relation to CAP analysis, Cifer and colleagues pointed out that long temporal autocorrelation of the (stationary) fMRI signal can explain the finding that a small fraction of time points suffices to approximate static FC (Cifer *et al*., 2017). More recently, Novelli and Razi asked whether edge-centric FC (Faskowitz et al., 2020; Zamani Esfahlani et al., 2020), a recently developed point-process method similar to CAP analysis performed in the connectivity space, can capture the non-stationarity of FC in resting brain activity. They mathematically showed that the results obtained with the edge FC method, at least in its present form, can be explained by assuming a stationary Gaussian distribution with a covariance matrix matched to real data. In the present study, we extended these previous studies by showing that the results obtained with CAP analysis, another widely used analysis technique for assessing resting brain activity, were similar for real fMRI data and for simulated data generated using the Laumann null model. This was surprising because the fact that a set of CAPs made from the time course of a single Seed ROI can be clustered into spatially heterogeneous modules appeals to our intuition that these modules represent distinct states of resting brain activity. The present results suggest that this intuition is incorrect. Spatially heterogeneous modules of CAPs were found with simulated data generated by a stationary null model. Moreover, many of the modules found in the simulated data were similar to the modules found in the real data. Thus, the presence of spatially heterogeneous modules of CAPs is insufficient to determine whether the data was generated by a mixture of distributions (*i*.*e*., non-stationary activity with multiple states). In Supplementary Figure 4, we replicated the findings of the original CAP study by Liu and Duyn (Liu and Duyn, 2013) in which CAPs obtained from surrogate data were spatially uniform. However, because these surrogate data broke down the static statistical properties of the data (e.g., covariance across voxels), the result cannot be attributed uniquely to the stationarity of the resting-brain activity. These studies collectively suggest that extra care needs to be taken when interpreting the results from these analysis techniques designed to extract dynamic structures of resting brain activity.

The assessment of non-stationary dynamics requires a temporal reshuffling approach or fitting an AR with non-zero delay (Liégeois *et al*., 2017). Recent studies have shown that fitting an AR is a promising approach to take into account the dynamic aspect of resting-brain activity (Liégeois et al., 2019). However, AR does not provide intuitive understanding of resting-brain activity, which could be important for guiding subsequent investigations. Thus, we believe that unification of an intuitive approach such as CAP with temporal reshuffling or AR would be an important topic for future studies of resting-brain activity.

We would like to emphasize that we are not claiming or trying to prove that resting brain activity is best represented by a stationary Gaussian distribution. In fact, careful statistical analyses suggest that resting brain activity is non-stationary (Liégeois *et al*., 2017). Similar to a previous study that examined sliding window FC analysis (Laumann *et al*., 2016), our intention was to make clear what can and cannot be concluded from CAP analysis. We would also like to note that the aim of the present study was not to deny the potential clinical usefulness of CAP analysis. Several studies have applied CAP analysis to clinical data and found valuable features of brain activity that characterize patient groups (Liu *et al*., 2018; Marshall et al., 2020; Rey et al., 2021; Yang et al., 2021). It is important to emphasize that the usefulness or clinical relevance of these features are not diminished by the present results (Liegeois R., 2021). Nevertheless, the present study sets a limit on how these features might be interpreted. For example, even when spatial patterns of modules are informative for distinguishing between CAPs from patients and healthy controls, it may be incorrect to interpret the result as evidence of distinct “meta-states” between the two groups. We believe that the correct interpretation of CAP analysis and sliding window FC analysis is indispensable for constructing realistic models of resting activity among healthy people and those affected by mental disorders (Wang and Krystal, 2014). Beyond fMRI studies, the present results indicate that appropriate surrogate data is likely to be important also in other settings such as micro-state analysis of electroencephalogram data (Michel and Koenig, 2018).

## Acknowledgements

We thank T. Ichikawa and M. Taki for discussion. Data were provided in part by the Human Connectome Project, WU-Minn Consortium (Principal Investigators: David Van Essen and Kamil Ugurbil; 1U54MH091657) funded by the 16 NIH Institutes and Centers that support the NIH Blueprint for Neuroscience Research; and by the McDonnell Center for Systems Neuroscience at Washington University. This study was supported by JSPS Kakenhi (20H05052 and 21H0516513 to TM, 19K20390 to TQP, 19H04914 and 20K07727 to KJ, 21H02806 to JC); a grant from Japan Agency for Medical Research and Development (AMED) to JC (grant number JP21dm0207086). a grant from Brain/MINDS Beyond (AMED) to TM (grant number JP20dm0307031); a grant from JST-PRESTO to TM; a grant from Narishige Neuroscience Research Foundation to TM.

## Supplementary Figures

**Supplementary Figure 1.**
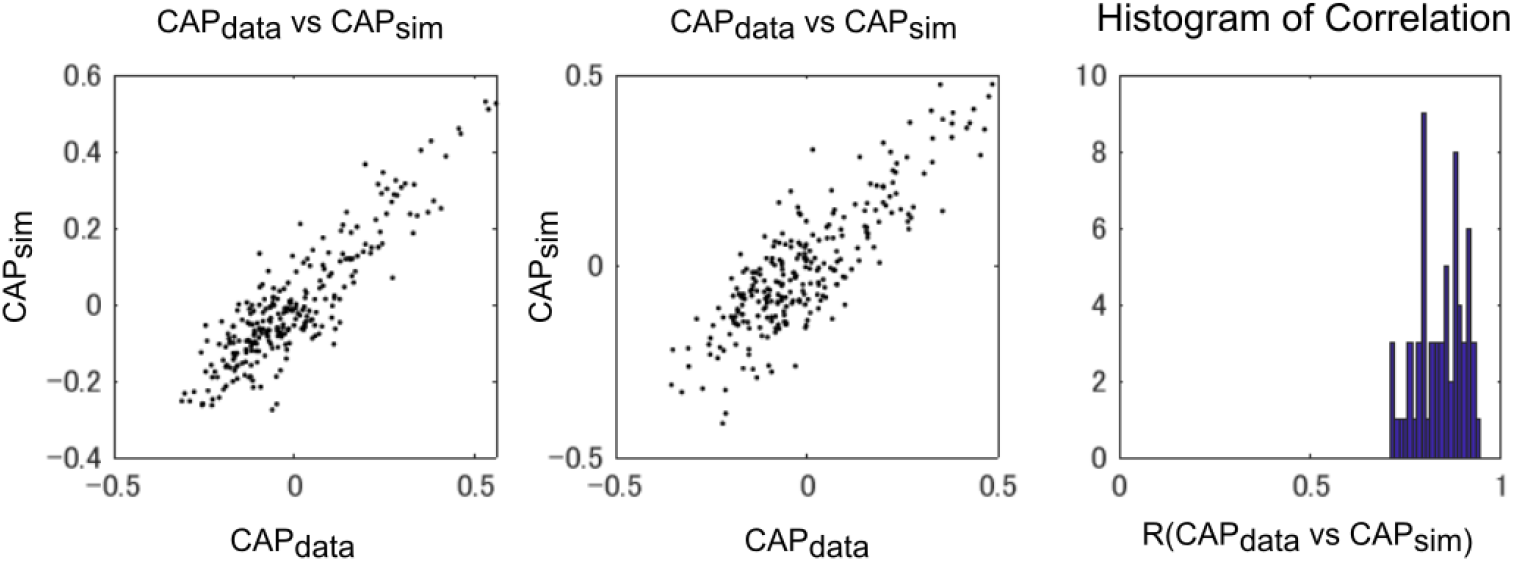
Panels on left and middle show comparison of mean CAPs obtained with real and simulated data for two subjects. Correlation coefficients are 0.92 and 0.88 for left and middle panels, respectively. Right panel show histogram of correlation coefficient between the mean CAPs for all subjects (mean ± SD, 0.84 ± 0.062; n = 64 subjects).

**Supplementary Figure 2:**
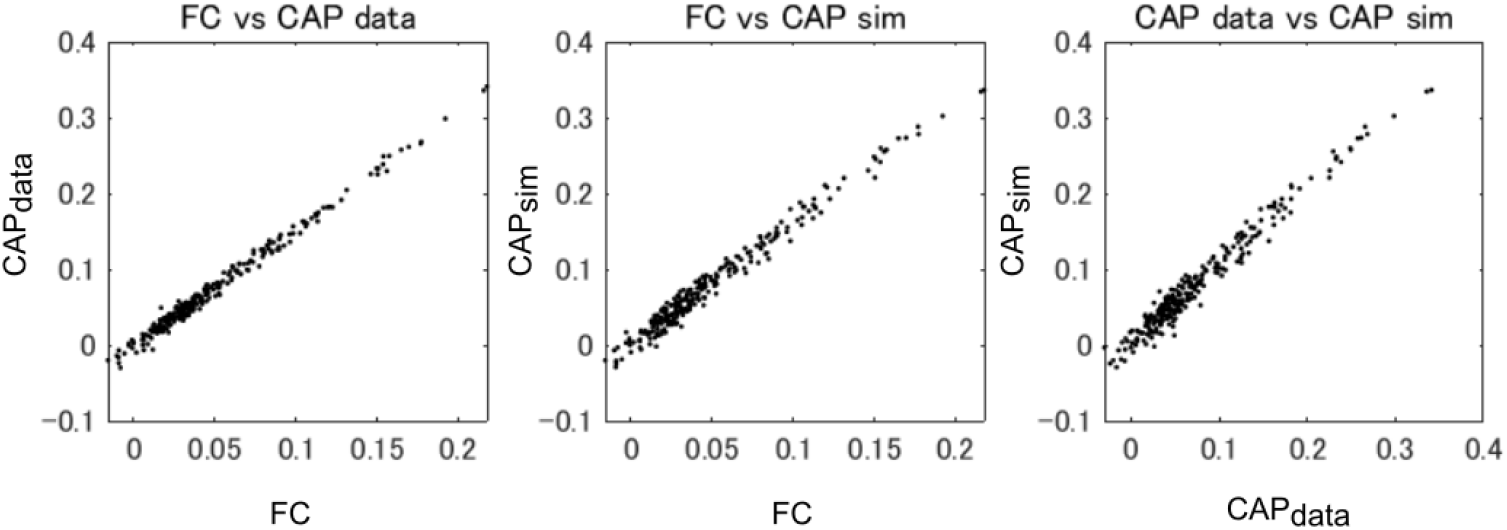
Same as Figure 1a-c but with 5% threshold for CAP detection.

**Supplementary Figure 3:**
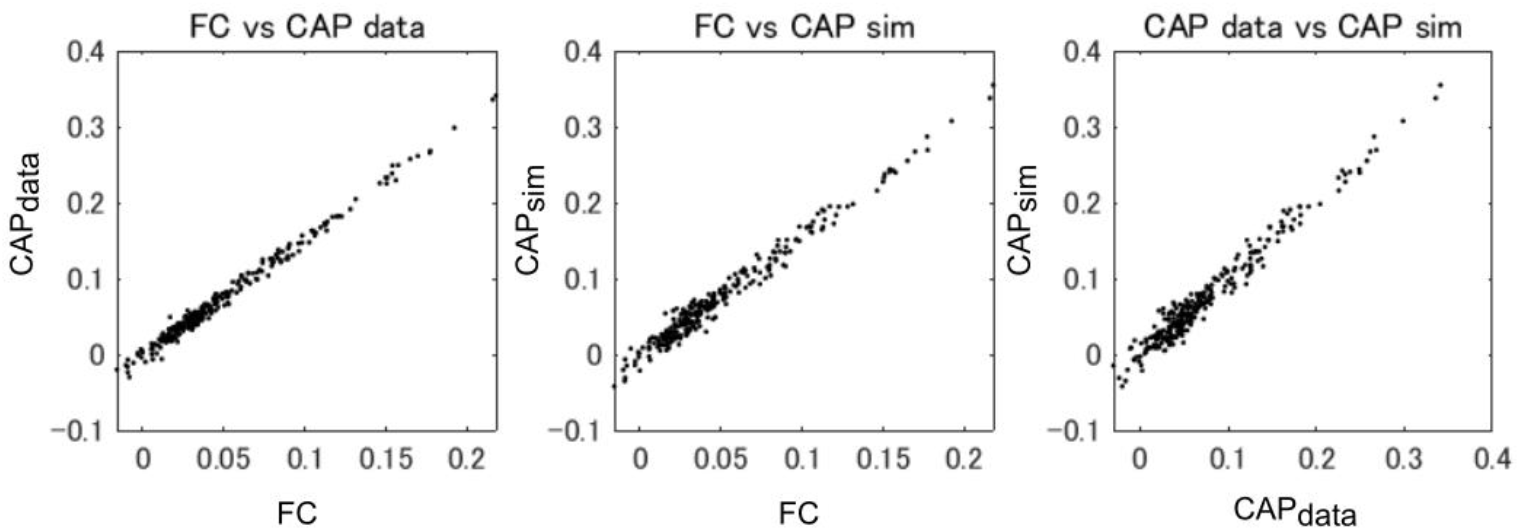
Same as Figure 1a-c but without GSR

**Supplementary Figure 4:**
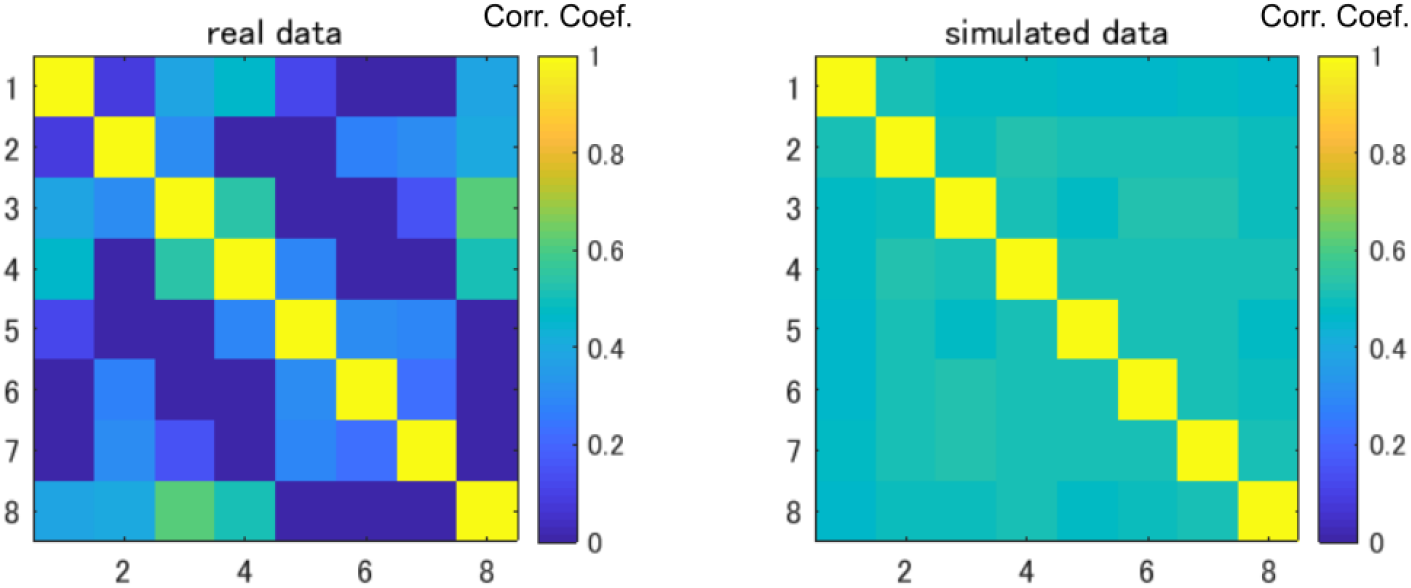
Same as Figure 2 but with simulated data with the Liu & Duyn-style null model (right panel). Compared with the real data (left panel) and the Laumann style null model (Figure 2, right panel), modules found with the Liu & Duyn-style null model were similar to each other (*i*.*e*., high off diagonal values).

**Supplementary Figure 5:**
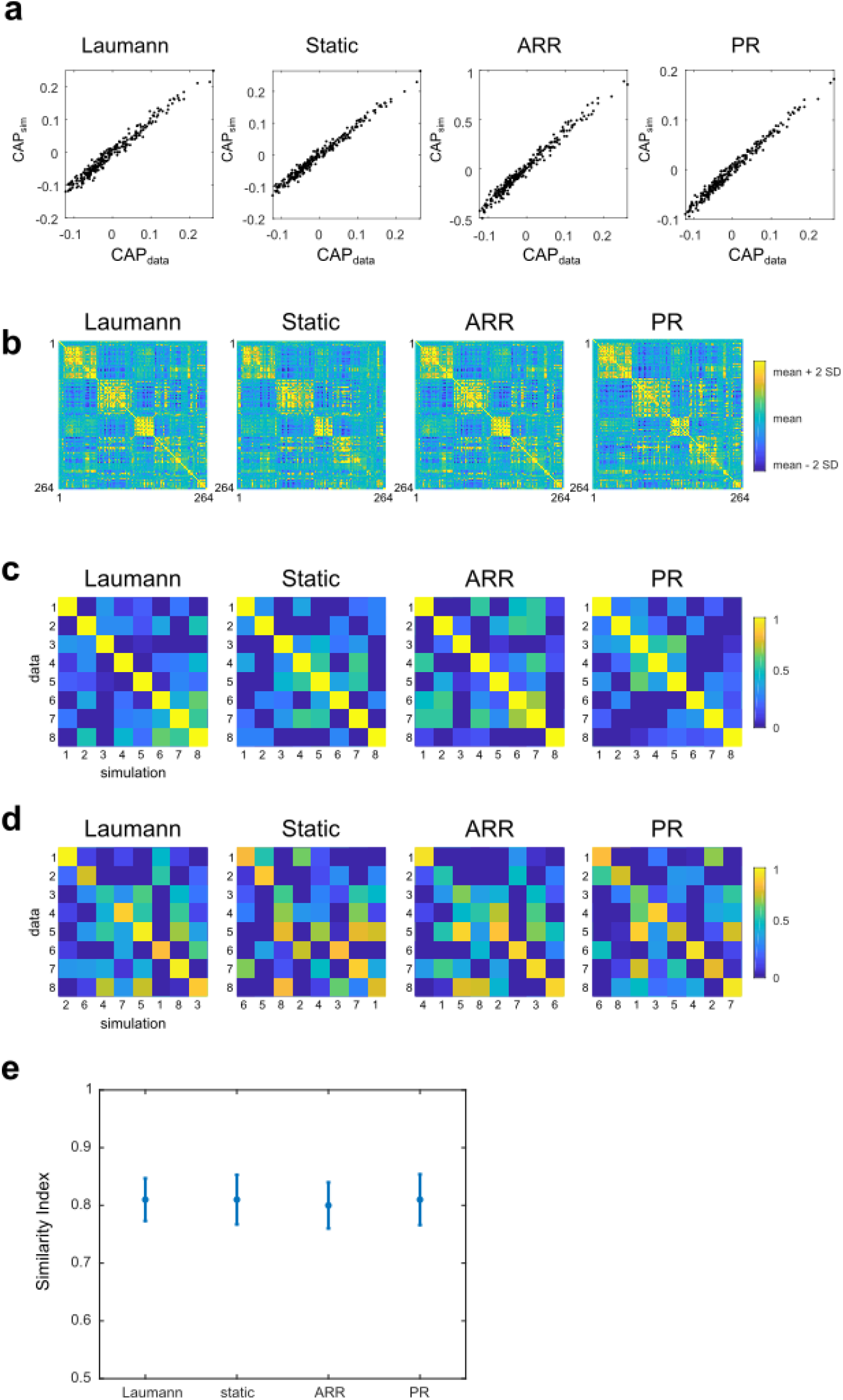
Panels a-e shows the results corresponding to Figure 1c&f, Figure 2b and Figure 3a but for all four null models. Similar results were obtained for all the null models, suggesting that the spatial structure of CAPs is determined by the covariance structure of the real data (which is the only factor retained in the static null). Panel e shows Similarity Index (same index as in Figure 3d) for four null models. Mean values across 264 ROIs are shown. Error bars indicate S.D.

**Supplementary Figure 6:**
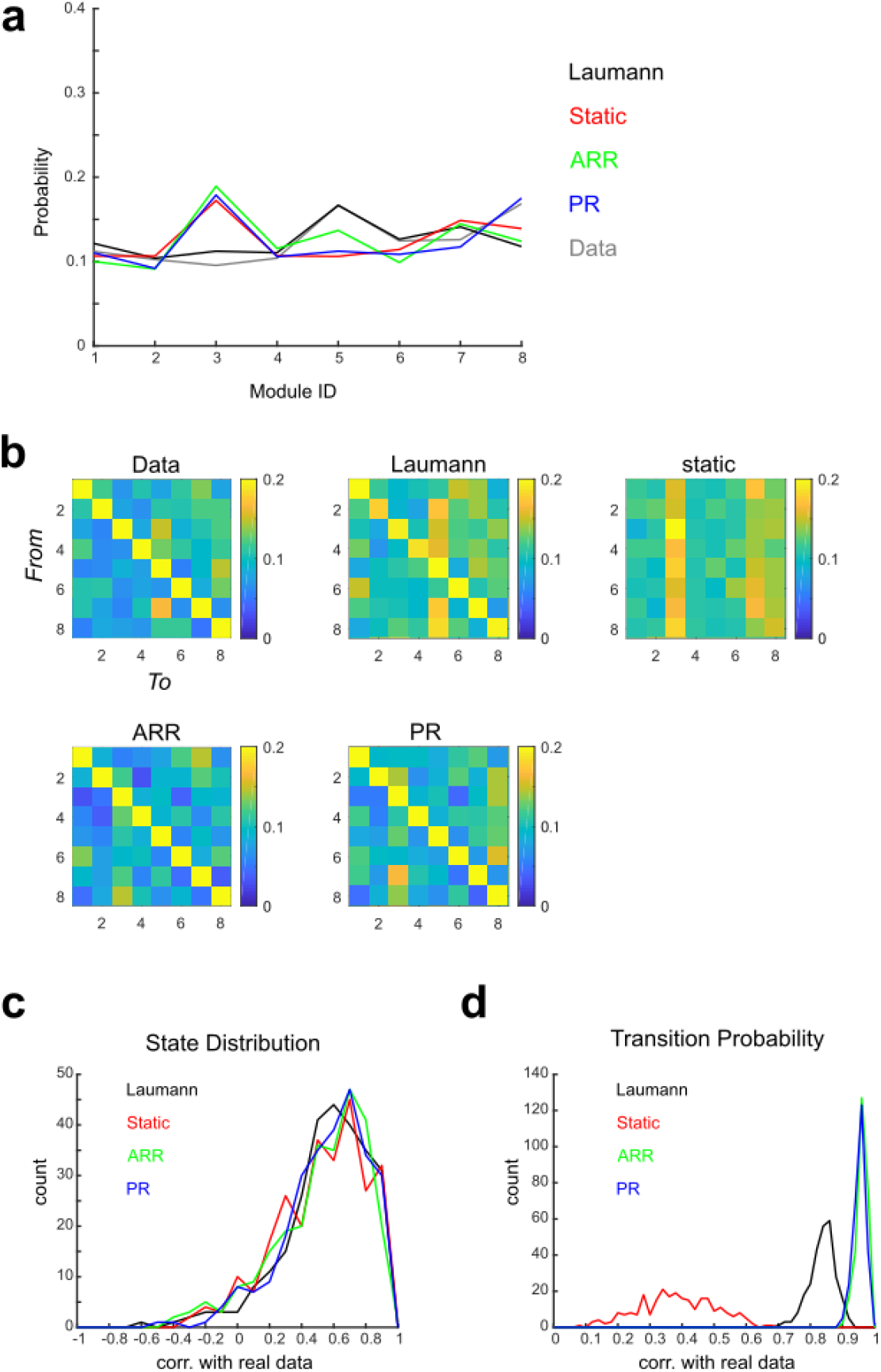
Distribution of states and Transition Probability Matrix. **a**. Examples of state distribution for the real data and four null models. **b**. Examples of transition probability matrices. **c**. Histogram of the correlation of state distributions for the real and a null model. **d**. Histogram of the correlation of transition probability matrices for the real and a null model.

